# The core microbiome of cultured Pacific oyster spat develops with age but not mortality

**DOI:** 10.1101/2023.10.27.564467

**Authors:** Anna Cho, Jan F. Finke, Kevin X. Zhong, Amy M. Chan, Rob Saunders, Angela Schulze, Snehal Warne, Kristina M. Miller, Curtis A. Suttle

## Abstract

The Pacific oyster (*Magallana gigas*, also known as *Crassostrea gigas*) is the most widely cultured shellfish worldwide, but production has been affected by mortality events. This includes mortality events in hatcheries that can threaten the seed supply for growers. There are several pathogens that cause disease in oysters, but in many cases mortality events cannot be attributed to a single agent, and appear to be multifactorial and involve a combination of environmental variables, microbial interactions and disbiosis. In many organisms, a mature microbiome provides resilience against pathogens and environmental stressors. In this study we investigated the microbiomes of cohorts of freshly settled oyster spat, some of which experienced notable mortality. Deep sequencing of 16S rRNA gene fragments did not show a significant difference among the microbiomes of cohorts experiencing different levels of mortality, but revealed a characteristic core microbiome with 74 taxa. Irrespective of mortality, the spat core microbiomes changed significantly in the relative abundance of taxa as the spat aged; yet, remained distinct from the microbial community in the surrounding water. The core microbiome was dominated by bacteria in the families *Rhodobacteraceae*, *Nitrosomonadaceae*, *Flavobacteriaceae, Pirellulaeceae* and *Saprospiraceae*. Of these, 14 taxa were indicative for the change in the core microbiome, which we designated as the “Hard-Core Microbiome”. The variability in diversity and richess of members of the core taxa decreased with oyster-spat aging, implying niche occupation. The study further accounts for the exchange of microbes with surrounding water during the core microbiome development. The observed shifts in the core microbiome with ageing oyster spat implies a crucial developmental period for the core microbiome of rearing spat.

**Importance:** The Pacific oyster (*Magallana gigas*, also known as *Crassostrea gigas*) is the most widely cultivated shellfish and is important to the economy of many coastal communities. However, high mortality of spat during the first few days following metamorphosis can affect the seed supply to oyster growers. Here, we show that the microbiome composition of recently settled oyster spat experiencing low or high mortality were not significantly different. Instead, development of the core microbiome were associated with spat aging and was partially driven by dispersal through the water. These findings imply the importance of early stage rearing conditons for spat microbiome development in aquaculture facilities. Furthermore, shellfish growers could gain information about the developmental state of the oyster spat microbiome by assessing key taxa. Additionally, the study provides a baseline micriobiome for future hypothesis testing on developing spat.

## Introduction

Native to Japan, the Pacific oyster (*Magallana gigas*, also known as *Crassostrea gigas*) is the most widely cultivated oyster worldwide (1). Despite its resilience to a broad range of environmental conditions, mass mortality events have been reported globally, including France (2), Australia (3), California, USA (4) and British Columbia, Canada (5). Mortality events occur at all life stages in both natural-spawning reefs (6) and farms (7–10). However, mortality incidents at the post-metamorphosis stage (spat) in hatcheries are of special concern as they can limit the availability seed to growers. Consequently, growers in the Pacific Northwest (11, 12) and Alaska (13) that rely heavily on hatchery spat have faced economic challenges. Despite numerous studies (2–5, 7–10, 14, 15), the causes of many mortality events remain elusive.

Studies have attributed the effects of ocean acidification (15, 16), temperature (17), farming location and practices (5), and emerging pathogens (18) as potential causes of mortality. More recent studies have shown that Pacific Oyster Mortality Syndrome (POMS), prevalent in adult oysters, is a polymicrobial disease involving a myriad of environmental factors (19), oyster genetics and multiple biotic agents such as Ostreoid Herpes Virus (OsHV) (14, 20, 21) and opportunistic bacteria (22).

There is evidence that the emergence of disease can be linked to changes in the host core microbiome (23–27). These reports lead to our initial hypothesis that the microbiomes between moribund (high mortality) and healthy (low mortality) spat are different, and our aim to establish a “core” microbiome of oyster spat.

The functional influence of a core microbiome on its host can be direct, such as for symbionts directly involved in metabolic processing (28), or indirect, as for transient core microbes that are present at one stage of a life cycle or under particular environmental conditions (29, 30). Core microbiomes can differ among tissues, life stages, species, and growth environments. There is evidence from corals and shellfish that a disturbed microbiome is associated with the host being susceptible to pathogens (25–27), including some which switch from a commensal or transient lifestyle. The state at which a disturbed microbiome becomes the cause of disease has been described as a pathobiome (23, 24).

These reports led us to hypothesize that there is a “core” microbiome that would be altered in spat experiencing high mortality.

The core microbiome of oysters has been characterized from different life stages, tissues, farm locations and broodstock using a variety of sequencing methods (17, 31–37). However, most studies have focused on microbiome dynamics in adult oysters under specific stressors (17, 32, 34) or candidate pathogens (8, 14, 21, 38, 39), and have not considered spat. However, the development of the spat microbiome is likely critical as they transform from a planktonic to a benthic lifestyle. This was demonstrated in an experiment in which immune genes and epigenetic expressions differed between mature larvae exposed to enriched microbes and ones exposed to hatchery-standard UV-treated seawater (40). Furthermore, the larvae with early microbial exposure had enhanced surivival when challenged with OsHV-1. Similarly, age-related (41) and metamorphosis-related microbiome shifts in insects (42, 43) amphibians (29) and fish (44) have been linked to changes in metabolic processes and defense against pathogens.

In the present study, the microbiomes of early-stage oyster spat were analyzed over 43 days, during which cohorts experienced different rates of mortality. We observed a core microbiome in spat that was distinct from that of the surrounding water and significantly changed in the relative abundance of taxa as the spat aged. At the same time, the variability in richness and diversity of members of the core microbiome decreased. However, the composition of the spat microbiome did not correlate significantly with mortality. These results demonstrate that the core microbiome of freshly settled Pacific Oyster spat changes with age, but is resilient to mortality.

## Results

### Data overview

A total of 47 samples, comprised of 39 oyster spat samples from nine cohorts, and eight water samples were collected from May to October 2014. Spat cohorts were reared at temperature, salinity and pH adjusted conditions (Table 1) according to industry standards. The age of spat ranged from zero to 43 days post-metamorphosis, some cohorts experienced up to 99% mortality within the first two weeks of the spat setting in the nursery tanks. As it is usual to have some mortality in developing spat, we defined ‘high mortality’ cohorts as ones that experienced > 50% mortality and ‘low mortality’ as ones with < 50% mortality.

**Table 1:** Growth parameters and mortality status of oyster spat samples. Cohort number and age in days post settling of spat, observed cohort mortality rates (Mort.), temperature (Temp.), salinity (Sal.), alkalinity (Alk.) and pH as measured in tanks.

| Sample Cohort | Age | Mort. | Temp. (°C) | Sal. ‰ | Alk. | pH |
| --- | --- | --- | --- | --- | --- | --- |
| BC01 | 5 | low | 17.8 | 31 | 3.12 | 8.52 |
| BC01 | 12 | low | 16.4 | 30 | 4.16 | 8.46 |
| BC01 | 19 | low | 20.2 | 25 | 3.76 | 8.37 |
| BC01 | 26 | low | 18.1 | 24 | 3.55 | 8.14 |
| BC05 | 4 | high | 21.8 | 22 | 4.25 | 8.08 |
| BC05 | 11 | high | 23.3 | 25 | 4.27 | 8.16 |
| BC05 | 17 | high | 19.4 | 22 | 4.2 | 8.58 |
| BC13 | 4 | high | 21.8 | 22 | 4.25 | 8.08 |
| BC13 | 11 | high | 23.3 | 25 | 4.27 | 8.16 |
| BC13 | 18 | high | 19.4 | 21 | 4.2 | 8.58 |
| BC13 | 25 | high | 24.3 | 25 | 2.74 | 8.09 |
| BC14 | 7 | low | 18.1 | 22 | 4.17 | 8.4 |
| BC14 | 14 | low | 24 | 25 | 2.92 | 8.12 |
| BC14 | 21 | low | 21.1 | 24 | 3.71 | 8.25 |
| BC14 | 28 | low | 18.7 | 26 | 3.87 | 8.17 |
| BC14 | 35 | low | 20.6 | 25 | 6.26 | 8.34 |
| BC15 | 3 | low | 24.4 | 23 | 2.78 | 8.19 |
| BC15 | 10 | low | 20.9 | 25 | 3.44 | 8.4 |
| BC15 | 17 | low | 19.2 | 25 | 3.21 | 8.59 |
| BC16 | 0 | high | 20.1 | 22 | 3.34 | 8.33 |
| BC16 | 7 | high | 18 | 26 | 3.3 | 8.2 |
| BC16 | 14 | high | 20.9 | 26 | 6.34 | 8.48 |
| BC18 | 1 | low | 15.4 | 26 | 3.44 | 8.47 |
| BC18 | 8 | low | 21.3 | 26 | 3.12 | 8.46 |
| BC18 | 15 | low | 13.4 | 29 | 3.15 | 8.78 |
| BC18 | 22 | low | 13.6 | 29 | 3.45 | 8.7 |
| BC18 | 29 | low | 12.5 | 30 | 3.31 | 8.76 |
| BC18 | 36 | low | 13.1 | 30 | 3.02 | 8.01 |
| BC18 | 43 | low | 11.3 | 30 | 2.89 | 7.98 |
| BC19 | 5 | high | 21 | 26 | 3.08 | 8.42 |
| BC19 | 12 | high | 13.4 | 29 | 3.15 | 8.78 |
| BC19 | 19 | high | 13.6 | 29 | 3.45 | 8.7 |
| BC19 | 26 | high | 12.5 | 30 | 3.31 | 8.76 |
| BC19 | 33 | high | 13.1 | 30 | 3.02 | 8.01 |
| BC19 | 40 | high | 11.3 | 30 | 2.89 | 7.98 |
| BC20 | 0 | low | 12.5 | 29 | 3.41 | 8.89 |
| BC20 | 7 | low | 12.4 | 30 | 3.28 | 8.85 |
| BC20 | 14 | low | 13.1 | 30 | 3.02 | 8.1 |
| BC20 | 21 | low | 11.3 | 30 | 2.89 | 7.98 |

Sequencing of 16S rRNA gene V4-V5 amplicons yielded 1,373,696 reads. The combined oyster and water samples produced 21,506 biological sequence variants, which were analyzed as unclustered OTUs (Operational Taxonomic Units). On average 27,450 reads per sample were generated. The reads were mapped to the OTUs to produced a frequency matrix and were rarefied to the minimum number of reads per sample (16,881). Taxonomic annotation based on SILVA (v.132) resulted in 91.8 % of OTUs being annotated to phylum, 87.3 % to class, 77.8 % to order, 73.1 % to family, 39.5 % to genus, and 13.2 % to species. Collapsing taxa at the family level resulted in 257 families, with *Rhodobacteraceae* (*Pseudomonadota*) and *Flavobacteriaceae*, *Nitrosomonadaceae* and *Saprospiraceae* (*Bacteroidota*)*, Rubinisphaeracea* and *Pirellulaeceae* (*Planctomycetota*) being dominant. There were no clear differences between water and oyster samples in the relative abundances of the top-ten bacterial families (Fig. 1). Other abundant families (data not shown) included *Cyanobiaceae (Cyanobacteria)* and *Vibrionaceae (Pseudomonadota).* Also, there was no significant difference in diversity between the microbiomes of ‘low mortality’ and ‘high mortality’ cohorts (Supplementary Table 1). A constrained distance-based redundancy analysis (dbRDA) showed weak separation between microbiomes of low-and high-mortality samples (Supplementary Figure 1), but mortality only explained 2.9% of the microbiome variation and was not significant (P-value 0.279).

**Figure 1:**
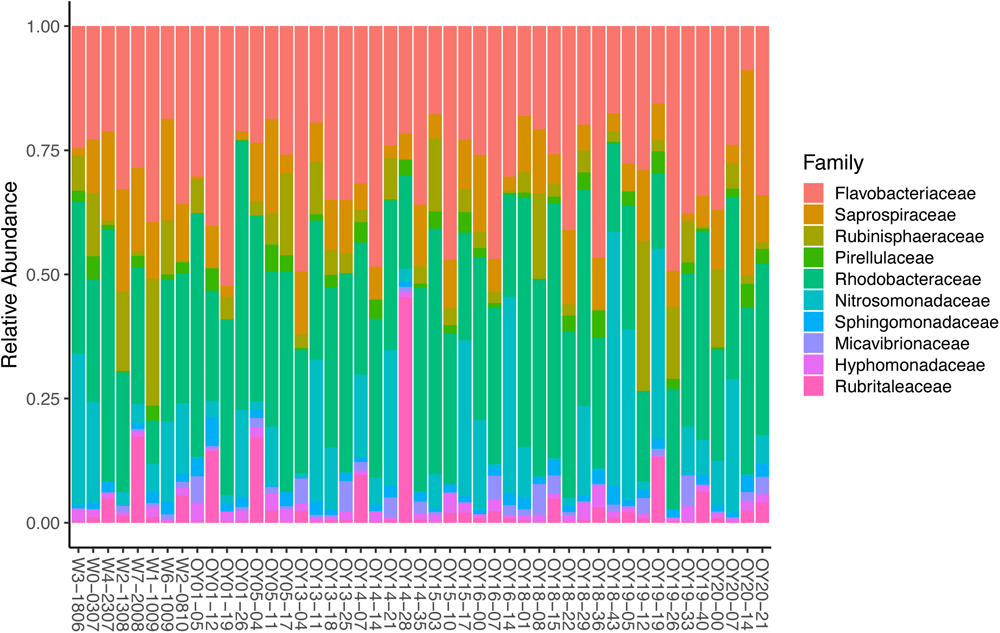
Overview of the relative abundances of the top ten bacteria families in the combined sequencing data based on 16S rRNA gene sequences. Water samples are labeled “W” followed by tank number and sample date (ddmm). Spat samples are labeled “OY” followed by cohort number, and the number of days post-settlement.

To assess the dissimilarity of microbial communities between oyster and water samples for all taxa, a constrained dbRDA of oyster samples against the sample source (i.e. tank water) was performed. The dbRDA (Figure 2) shows that water samples (blue triangles) were tightly clustered and clearly separated from oyster samples (red circles). Although there was a lot of variation among the oyster samples, they were significantly different from the water samples (P-value <0.001), based on a PERMANOVA.

**Figure 2:**
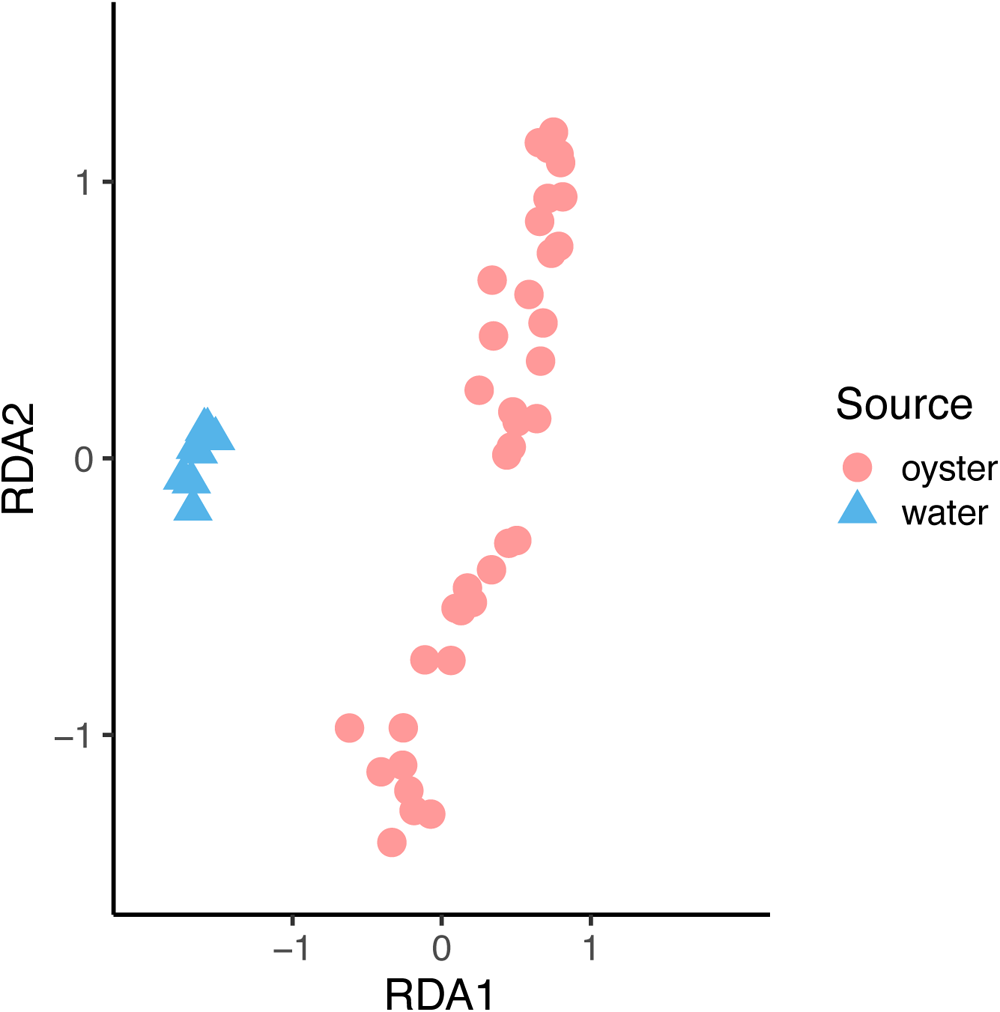
Constrained dbRDA of the community composition of water and oyster samples against the sample source. Blue triangles are water samples; red circles are oyster samples.

### The core microbiome of Pacific Oyster spat

To identify taxa constituting the core microbiome of spat, we selected OTUs present in more than 50% of all oyster samples, and that were differentially represented in oysters compared to water. This approach identified 74 core OTUs that were predominantly in the families *Rhodobacteraceae, Flavobacteriaceae*, *Nitrosomonadaceae*, but also included *Saprospiraceae*, *Haliaceae*, *Bdellovibrionaceae* and unassigned OTUs (Summarized in Supplementary Table 2). An unconstrained dbRDA of the core microbiome composition in oyster samples showed a pattern of shifting community similarity with spat age (Figure 3). While some outliers are observed, there is a gradual transition of the core microbiome as the spat aged, which was confirmed to be significant in a constrained dbRDA and subsequent ANOVA (P-value <0.001) of core microbiome similarity against age. Notably, the dbRDA of the core taxa did not show a correlation between the core microbiome and mortality rates.

**Figure 3:**
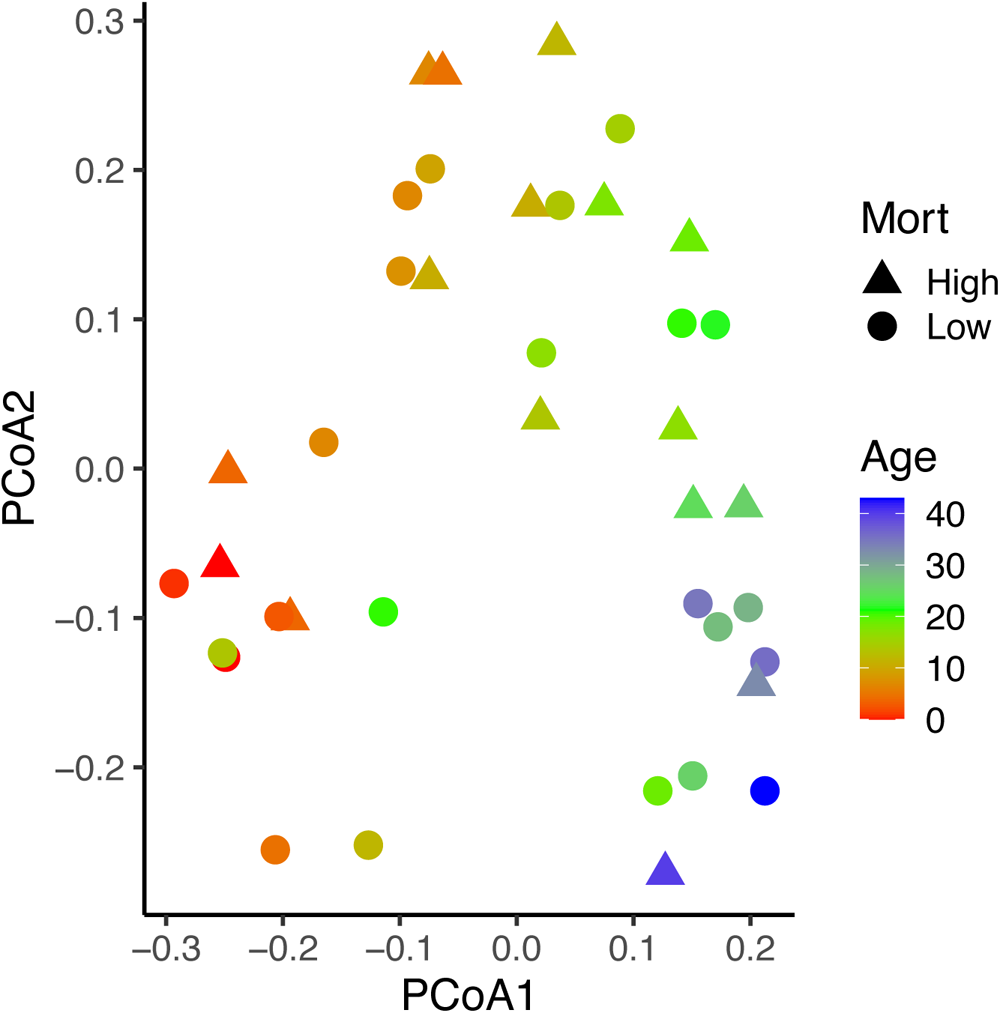
PCoA (unconstrained dbRDA) of the community composition of core OTUs. Shapes indicate whether cohorts experienced low <50% /d (circles) or high >50% /d (triangles) mortality. Colour indicates the age in days post settlement.

To identify representative OTUs from the core microbiome that best describe the age-related microbiome shift, a Balance-Tree Analysis was used. The analysis calculates balance values for samples based on the difference in ratio of selected OTUs with regard to age. The analysis identified five OTUs belonging to taxa within the *Rhodobacteraceae, Flavobacteriaceae* and *Nitrosomonadaceae*, which were used to calculate balance values for each sample (Supplementary Table 3). The combination of the selected OTUs resulted in a significant linear correlation (Supplementary Figure 2) between their relative abundance in each sample and age (R^2^ =0.9266; P-value<2e-16).

### Evolution of the core microbiome with age

To better understand how the core microbiome in oyster spat changed with age, spat samples were categorized into six age classes (days post-metamorphosis), A (0 to 4), B (5 to 11), C (12 to 17), D (18 to 23), E (24 to 29), F (>=30), which had respective sample sizes (n) of 6, 9, 8, 6, 5 and 5. Generally, the richness of the core microbiome increased with increasing age, more so in age classes A to C than D to F. Meanwhile the variability in richness within age classes decreased (Figure 4). Similarly, alpha diversity increased from A to C, while average diversity and especially the variability in diversity dropped in classes D to F. For the six age classes, corresponding beta diversities were 0.87, 1.51, 1.43, 1.33, 1.22 and 1.19, while gamma diversities were 2.17, 3.26, 3.33, 3.13, 2.84 and 2.65, respectively. The frequency of core OTUs across samples in each age class mirrored the diversity indices (Figure 5). Most core OTUs occurred less frequently across samples in age classes A to C, but were more prevalent in D, E and F. However, some OTUs showed the opposite trend; two OTUs in the *Rhodobacteraceae* had a high frequency in A, but were almost absent in F. Overall, eleven OTUs, mainly in the *Nitrosonomadaceae* and *Rhodobacteraceae*) showed a significant increase and two OTUs (*Rhodobacteraceae*) showed a significant decrease in linear models with progressing age class (Supplementary Table 3).

**Figure 4:**
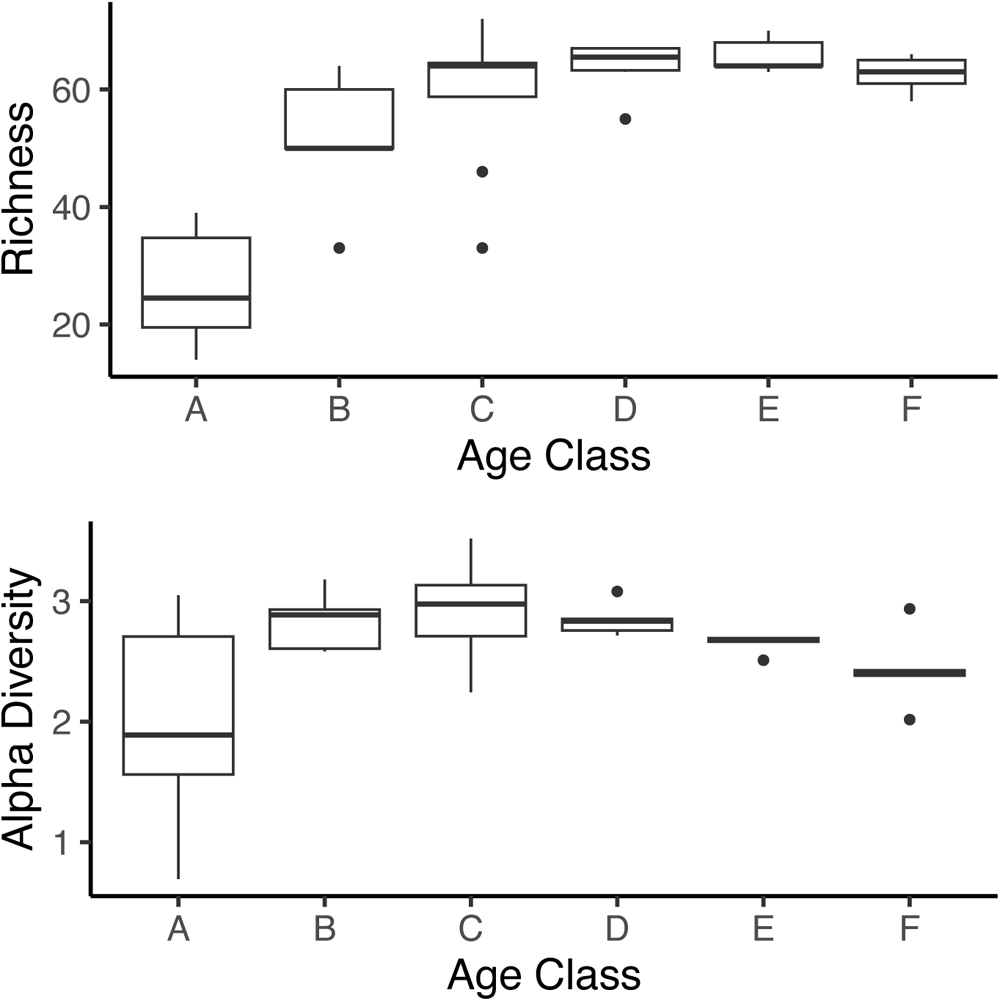
Core microbiome diversity indices for spat age classes, indices were calculated based on the community compositions of samples per age class. Top graph depicts OTU richness; bottom graph shows alpha diversity, age classes (days post settlement), A (0 to 4), B (5 to 11), C (12 to 17), D (18 to 23), E (24 to 39), F (>=30), which had respective sample sizes (n) of 6, 9, 8, 6, 5 and 5. Box plots describe the minimum and maximum values, lower and upper quartiles and median, outliers.

**Figure 5:**
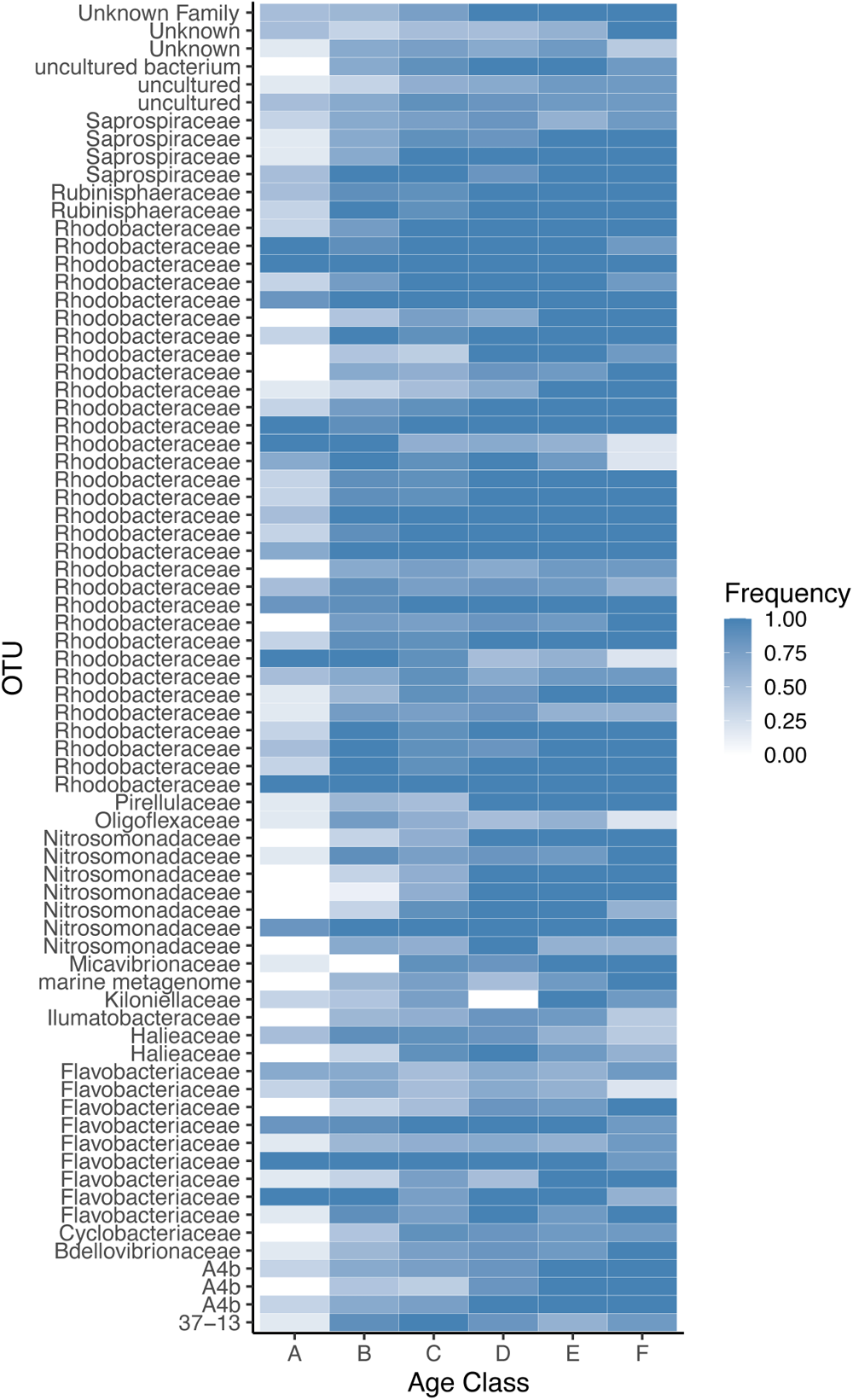
Heat map of the frequency of core OTUs across samples per age class. Colour tone describes frequency per age class; white is absence and dark blue is presence. OTUs are defined at the genus level, but family annotation is labeled.

Comparing early age classes A to C against late age classes D to F in an Indicator Species Analysis (45) uncovered eight significant indicator species. Two OTUs in the *Rhodobacteraceae* were indicators for the early age classes, while three OTUs in the *Nitrosonomadaceae*, and one each in the *Rhodobacteraceae, Pirellulaceae* and A4b (phylum *Chloroflexi*) were indicator species for the later age classes (Supplementary table 3). The combined analyses were consistent for 14 OTUs that had significant linear correlations to age. Several of these OTUs increased in frequency with age, were indicator taxa for the older age classes and were key taxa in the balance model. Accordingly, the OTUs that decreased in frequency with age were indicator taxa for younger age classes (Supplementary Table 3).

Finally, a Neutral Community Model (46) was applied to explore the dispersion of core OTUs from surrounding water to spat during progression of the core microbiome. This analysis could only be performed on a subset of 47 core OTUs that were present in spat and water samples. The predicted frequency of the 47 OTUs in the oyster communities was calculated based on their mean relative abundance in water samples. Twenty OTUs (grey circles) with observed frequencies within the 95% confidence interval of their predicted frequencies were designated to be neutrally dispersed from water to oyster spat (Figure 6). Ten OTUs with higher observed than predicted frequencies (red circles) were designated to be overrepresented in oysters, and 17 OTUs with lower observed frequencies than predicted (blue circles) were designated to be underrepresented. The neutrally dispersed OTUs were predominantly 11 OTUs in the *Rhodobacteraceae*, two in the *Saprospiraceae*, and one in each of the *Nitrosomonaceae* and *Flavobacteriaceae* (Summarized in Supplementary Table 4). Six of the overrepresented OTUs were in the *Rhodobacteracea*, while the underrepresented OTUs were predominantly in the *Flavobacteriaceae* and *Rhodobacteracea*.

**Figure 6:**
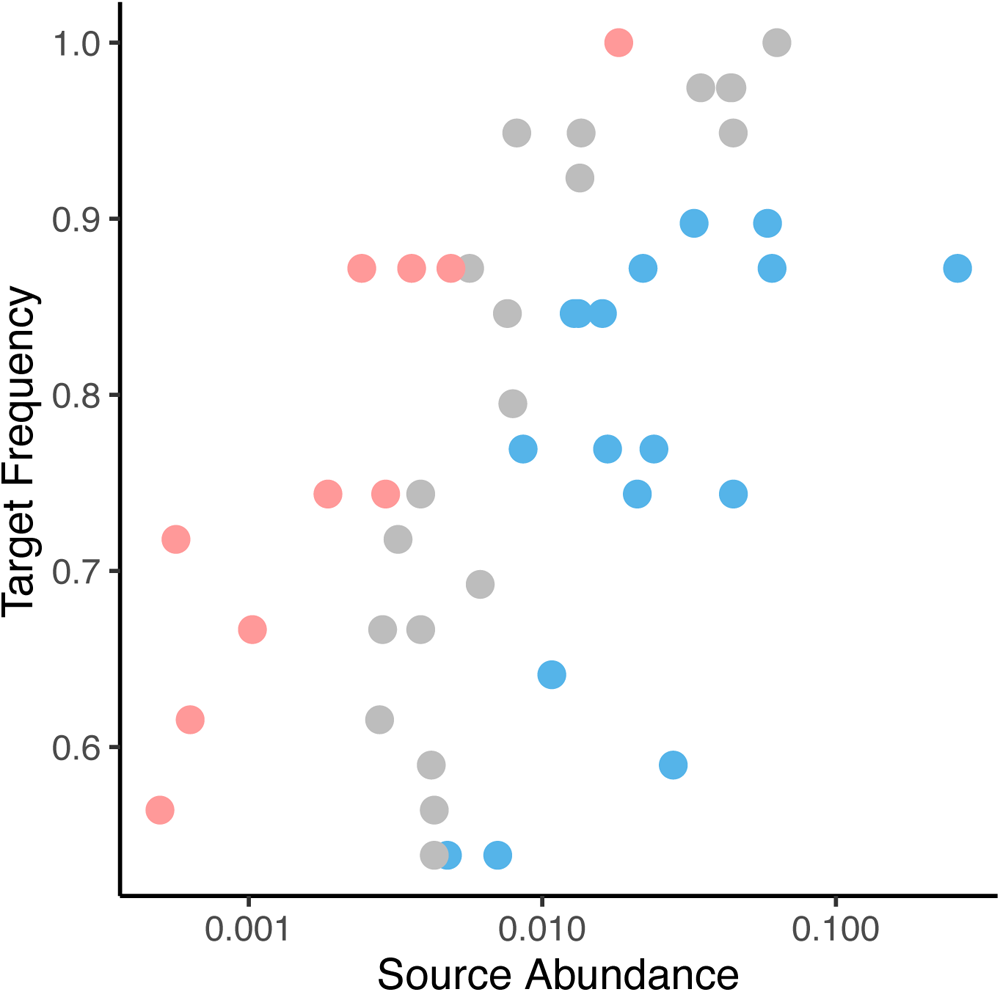
Mean abundance in water communities (Source Abundance) and frequency across oyster spat samples (Target Frequency) for OTUs included in the neutral community model analysis. Colours indicate the dispersion mode: red for OTUs overrepresented in oyster samples; blue for OTUs overrepresented in water samples; grey for OTUs where the observed frequency matched the predicted frequency in the neutral community model.

## Discussion

We explored changes in the microbiome of recently settled Pacific Oyster spat that had experienced a range of mortality. Samples were collected to reflect microbiome development in recently settled spat grown under standard aquaculture practices (11). The composition of the spat microbiome was not correlated with mortality; however, we revealed a core microbiome that differed significantly from that in the water. Moreover, the core microbiome was defined by a relatively small number of taxa that changed in relative abundance with age, and were dispersed through water. These observations and their implications are elaborated on below.

In total, 21,506 OTUs were identified, providing high taxonomic resolution; 73 % of these were classified at the family level for subsequent analysis. The most abundant OTUs were in the *Rhodobacteraceae, Flavobacteriaceae*, *Nitrosomonadaceae* and *Saprospiraceae*. Bacteria in these families are common in aquatic environments and belong to the major phyla *Bacteroidetes* and *Pseudomonadota* (*Proteobacteria*), which also occur in the hemolymph of adult and juvenile oysters (17, 32, 34, 47, 48), as well as in marine waters (49). Additionally, members of the *Planctomycetota* and *Cyanobacteria* were abundant, and are also commonly found in seawater (49). Overall, at the family level the dominant bacteria were consistently present across oyster spat and water samples. Nonetheless, given that the dbRDA and ANOVA analyses showed significant differences between the OTUs of the oyster and water microbiomes, supports using the OTU data from the water as a comparator to identify the core microbes characteristic of oysters. Differences between the composition of the microbial communities in oysters and the surrounding water resembles observations from previous studies; yet, in contrast to King et al. (36) we did not find a significant correlation between microbiome community similarity and mortality.

While there is no single definition of a core microbiome, we followed the broad definition that the core microbiome is comprised of microbial taxa that are consistently associated with given ecological or biological habitats (50, 51). Consequently, the core microbiome was based on taxa that were more abundant in spat than in the adjacent water, and which had a frequency over 50% across spat samples. In a study on coral core microbiomes the number of taxa and their relative abundance drastically increased when the frequency cut-off values was lowered from 90% to 50% (52). This suggests that a higher percentage cut-off value would mask important shifts of the core microbiome over time. Since in our study the microbial composition was not significantly different between ‘low-mortality’ and ‘high-mortality’ spat samples, we decided to include all spat samples when defining the core microbiome. It should be noted that sample processing and sequencing methods can affect the interpretation of microbiome analysis (31).

Based on the described criteria, 74 out of 21,506 OTUs were selected as part of the core microbiome; primarily these OTUs belonged to the families *Rhodobacteraceae, Nitrosomonadaceae, Flavobacteriaceae*. These families match previously published work on mature oyster microbiomes over time (17, 48), differences can be explained by variations of body sites and sampling location that were described by Stevick et al. (48) and Dube et al. (47). While several families are similar the number of identified core taxa is relatively high in contrast to a study on the seaweed microbiome that identified only 24 core taxa in the *Rhodobacteraceae, Flavobacteriaceae* and *Saprospiraceae* for different host life stages (53).

In the present study the relative abundance of taxa in the core microbiome was significantly correlated to spat age (Figure 3). Similar shifts in microbiome composition with age have been documented in oysters from different life stages (33) and a recent study on the spat microbiome under experimental temperature and pCO2 stress also found a change of the microbiome composition with age (54). A correlation between the microbiome development and host age has also been documented in corals, insects and fish (41, 43, 55, 56). Notably, in a separate project on mature oyster microbiomes, no such correlation with age was apparent (37), highlighting the importance of the core microbiome development at early age. The shift of core microbiomes was further quantified by balance-tree analysis; sample age explained 93% of the occurrence of five reference OTUs in the *Rhodobacteraceae, Flavobacteriaceae* and *Nitrosomonadaceae*. Changes of the relative abundance of *Flavobacteriaceae* have also been induced by temperature and infection stress in oysters (17). Similarly, Trabal-Fernandez et al. (33) observed in oysters that at the genus level bacteria in the *Pseudomonadota* varied between post-larval and adult stages. The observed shifts in core microbiome composition within families can be explained by successful niche occupation of phylogenetically and ecologically similar taxa.

We also investigated the dynamics of temporal shifts in the core microbiome. For most core OTUs the frequency increased with progressing age classes, the eleven OTUs with significant increase in frequency may play a key role in oyster development. Similar patterns of microbiome changes with host development were observed in fish (44, 55, 56), insects (42, 43), but especially corals (41). Williams et al. (41) and Yan et al. (56) also observed that most OTUs increased in their relative abundance with early coral and fish microbiome development, while few OTUs were lost and replaced in older spat. The indicator species analysis further supported this observation with six OTUs that were characteristic for later age classes (D, E, F) and only two *Rhodobacteracea* OTUs that were indicator taxa for the early age classes (A, B, C). Furthermore, the core microbiome development was reflected by an initial increase of taxa richness and diversity with age, but subsequent decrease in the variability of core taxa. Again, Williams et al. (41) observed a similar increase in microbiome richness in developing corals, while Yan et al (56) showed decrease in alpha diversity of the gut microbiome during metamorphosis of fish larvae.

Finally, the neutral model analysis of core OTUs revealed the dispersal of taxa from rearing water to oyster microbiomes (46). The 20 *Rhodobacteraceae* OTUs fitting the neutral model were likely dispersed from water to the oysters as they aged. Of the OTUs not fitting the neutral model, the ten overrepresented OTUs appear to be selected for in the oysters, while 17 underrepresented OTUs appear to be selected against.

Dispersed and selected for OTUs are likely permanent members of the core microbiome, but the OTUs that are selected against are more likely to be lost from the core microbiome during aging. However, the OTUs that don’t fit the neutral model may also be the result of stochastic processes of dispersal and losses (57). These findings show that dispersal and selection processes are shaping the oyster core microbiome as has been seen in other marine environments (58). Furthermore, studies on coral microbiomes also suggest that microbes are dispersed through water (59), as well as vertically transmitted (60).

Altogether, bacteria in the families *Rhodobacteraceae, Flavobacteriaceae, Nitrosonomadaceae*, *Pirellulaceae* and the phylum *Chloroflexota* (family label A4B) have a significant role in the core microbiome development across analyses. The family *Rhodobacteraceae* comprises diverse groups of marine bacteria, from phototrophs and chemoheterotrophs involved in sulfur and carbon cycling, to symbionts of other marine organisms (61, 62). Previous studies also found that members of the *Rhodobacteraceae* can originate from algal feed stock in aquaculture (27). The families *Nitrosomonadaceae* (*Pseudomonadota*) and *Pirellulaceae* (*Planctomycetota*) comprise groups of ammonia oxidizing bacteria, and have been associated with deep-sea octocorals (*Paramuricea placomus*) (63) and kelp (*Macrocystis pyrifera*) (64). *Nitrosomonadaceae* have also been recorded in coral microbiomes (65, 66). These nitrifying taxa could be ubiquitous in marine filter-feeders and important in biogeochemical cycles (64, 67), or simply opportunists occupying ammonium-rich systems. The families *Flavobacteriaceae* and *Vibrionaceae* include common aerobic marine microbes, as well as putative pathogens (27, 68–70). Interestingly, the phylum *Chloroflexota* which encompasses a metabolically diverse group of bacteria has been reported in the gut of other adult oyster species (32, 71), and is one of the core taxa in sponge microbiomes (72), suggesting they may be a common taxon in filter-feeding marine animals. Furthermore, many of the core bacteria are dispersed throught the water column. Fourteen OTUs from these bacterial families appear to have an outsized role in the core microbiome development and could thus be considered the “Hard-Core Microbiome”. Assessing these key taxa of the core microbiome can provide shellfish growers with information about the developmental state of the oyster spat microbiome. The roles of the core taxa remain unclear, warranting research on the key metabolically relevant gene expressions with an emphasis on changing filter-feeding behaviour and physiology.

## Materials and Methods

### Oyster and Water Samples

Pacific oyster spat and water samples from a commercial oyster hatchery in British Columbia, Canada, were collected over several months in 2014. The broodstock was a collection of naturally spawning Pacific oysters from Discovery Passage, BC, Canada. Spat were grown by industry standards under optimal conditions at 21°C, pH 8.1-8.2, salinity 25 ‰ and 3.3 mg/L alkalinity; intake water was filtered through sand and 5-μm filters. Tank temperatures were measured using a mercury thermometer and salinity was measured with a refractometer. Water pH was measured with a glass probe pH meter (Jenco, San Diego, CA) and alkalinity was measured with a HI901 titrator (Hanna Instruments, Smithfield, RI). Spat size ranged between 400 and 1000 μm, and the percentage mortality were assessed daily using a microscope. Each week, about 15 to 30 spat were collected in 2-mL screw-cap cryovials for all. The percentage mortatliy ranged between 10% to 99% for each sampling time point; cohorts with >50% mortality were classified as “high mortality”, while <50% mortality was considered “low mortality”. Corresponding water samples were taken by filtering 2-L of tank water through 0.22-μm pore-size cellulose acetate membranes (Millipore Sigma, Burlington, MA). Samples were stored at −80°C until processing.

### DNA extraction

Approximately 20 oyster spat were transferred into sterile 1.5-mL microcentrifuge tubes using a sterile spatula or pipette. Using two 3.2-mm diameter chrome steel beads (Bio Spec Products Inc., Bartlesville, OK) the spat were homogenized in a Mixer Mill MM300 (Retsch GmbH, Haan, Germany) for 1 min with a frequency of 1/30 s. At least 2 mg of tissue homogenate from each tube was transferred to a new microcentrifuge tube, and DNA extracted using a DNeasy Blood & Tissue kit (Qiagen, Hilden, Germany) per manufacture’s protocol, with an added lysozyme (10 mg/mL) lysis step after Proteinase K addition. DNA from water samples was extracted using a DNeasy PowerWater kit (Qiagen, Hilden, Germany) following the manufacturer’s protocol. DNA extracts were quantified with the Qubit dsDNA HS Assay Kit (Life Technologies, Carlsbad, CA).

## 16S rRNA gene amplification, library preparation and sequencing

The 16S sequencing libraries were prepared following the Illumina MiSeq 16S Metagenomic Preparation Guide using the Nextera XT v2 Kit (Illumina, San Diego, CA), with the following minor modifications. PCR mixtures contained 1 to 10 ng of template DNA in a 25 μL reaction volume. The V4-V5 (515F-926R) hypervariable regions of the small subunit (SSU) 16S ribosomal RNA coding gene were targeted for amplification with barcoded forward and reverse primers. Conditions for PCR were denaturation for 5 min at 95°C followed by 34 cycles at 95°C for 45s, 50°C for 45s, 68°C for 90s and a final extension at 68°C for 10 min. The reaction mix contained 1X Q5 Reaction Buffer (New England Biolabs (NEB), Ipswich, MA), 0.4 μM barcoded forward and 0.4 μM reverse primers, 0.1 U Q5 High-Fidelity DNA Polymerase (NEB, Ipswich, MA), 0.8 mM dNTPs, 1X BSA (100x), 2.5 mM MgCl_2_. The Index-PCR reaction was performed in a 25 μL volume containing 10 ng DNA, using Nextera XT v2 Index 1 and 2 Primers. The reaction mix contained 1X Q5 Reaction Buffer (NEB, Ipswich, MA), 1.25 μL each of Nextera XT v2 Index 1 and 2 Primer, 0.1 U Q5 High-Fidelity DNA Pol (NEB, Ipswich, MA), 0.4 mM dNTPs, 2 mM MgCl_2_. Amplicon libraries were purified and size-selected using Agencourt AMPure XP Beads (Beckman Coulter Inc., Brea, CA), and quantified with the Qubit dsDNA HS Assay Kit (Life Technologies Inc., Carlsbad, CA). The Index attachment was verified by quantitative PCR (qPCR) using the SsoFast EvaGreen Supermix (Bio-Rad Laboratories Inc., Hercules, CA) and KAPA DNA Standards (KAPA Biosystems, Wilmington, MA) on a C10000 Touch PCR block with a CFX 96W Reaction Module (Bio-Rad Laboratories, Inc., Hercules, CA). Libraries were pooled, and gel-purified in a 1.5 % agarose gel (1x TBE) by excising the 560 base-pair (bp) band using a Zymoclean Gel DNA Recovery Kit (Zymo Research Corp., Irvine, CA), following the manufacturer’s instructions. The pooled libraries were validated with a Bioanalyzer High Sensitivity DNA chip (Agilent Technologies, Inc., Santa Clara, CA) and sequenced at the BRC Sequencing Core at The University of British Columbia on a MiSeq platform using 2 x 300 bp paired-end chemistry (Illumina, San Diego, CA).

### Bioinformatic and statistical analysis

Raw read data were demultiplexed followed by removal of primers, low quality reads (Phred < 29) and chimeras using Quantitative Insights into Microbial Ecology (QIIME 2) v 2018.6.0 (73). Forward and reverse reads were merged using the Divisive Amplicon Denoising Algorithm 2 (DADA2) in QIIME 2 (73, 74). The method produced a list of features, each representing a biological sequence variant, generated by the *de novo* error correction method of DADA2. Features were designated as operational taxonomic units (OTUs) based on the 99% sequence similary against SILVA reference database (release v.132) using QIIME2 (73, 75). A sequence similarity tree was generated using *de novo* multiple sequence alignment (MAFFT) (76), divergent OTUs with long branches were searched using the Basic Local Alignment Search Tool (BLAST) (77), and mitochondria, chloroplasts, metazoan and singletons were filtered from the OTU table. OTU frequency in samples was determined by read recruitment using QIIME2 (73), the frequency matrix was normalized to the lowest number of total reads per sample by rarefication with the VEGAN package (78) in the R environment (79) producing representative microbial communities per sample.

A taxonomic overview was generated by collapsing OTUs at the family level with the corresponding phyla. Microbial community composition was compared between water and oyster samples in a constrained distance-based redundancy analysis (dbRDA) using VEGAN; significance of the separation was assessed by a PERMANOVA test. Differential abundance of taxa between water and oyster samples was determined with the DESeq2 package (80) in R. To compare the similarity in community composition among samples the Bray-Curtis dissimilarity index was calculated and ordinated using an unconstrained dbRDA in VEGAN, with subsequent ANOVA test for significance. This is based on shared OTUs that were present in more than 50% of all oyster samples, and were differentially represented in oyster and water microbiomes. Taxa with the best linear correlation to age were determined with the Selbal package (81). Indicator taxa for age classes were identified with the Indicspecies package (82, 83). Dispersion of taxa from the water to oysters were tested with a neutral community model based on Sloan et al. (46) in R.

### Core microbiome

Similar to other studies, we defined the core microbiome as shared operational taxonomic units (OTUs) that occured in more than 50% of oyster samples, and which were differentially represented in spat than in the surrounding water (26, 50, 84). The core microbiome was defined based on spat that experienced low or high mortality as there was no significant difference in the microbiome composition between groups.

## Data Availability Statement

The sequencing data used in this study have been deposited in the NCBI Short Read Archive (SRA) under the accession number PRJNA1032956. All other data supporting the findings of this study are provided in the manuscript or as supplementary files.

## Acknowledgements

We thank Laura Parfrey and Andrew Loudon for advice on the neutral community model analysis. We also thank Corey Holt and Vittorio Boscaro for improvements to the manuscript. This project was supported by the Gordon and Betty Moore Foundation (Grant: GBMF #5600; to C.A.S and K.M.M.); the Natural Sciences and Engineering Research Council of Canada (Grants: RGPIN-2015-05896, RGPIN-2020-06519 to C.A.S.) and an infrastructure award from the Canada Foundation for Innovation and British Columbia Knowledge Development Fund (Project #25412 to C.A.S.). Hatchery collections were supported by Island Scallops Ltd., and an Aquaculture Collaborative Research and Development Program (ACRDP) grant from Fisheries and Oceans, Canada (Grant: P-14-02-001 to K.M.M). A.C. was supported by the Pacific Research Board (Grant: NPRB GSRA#1335) and an Ocean Leader Graduate Fellowship from The University of British Columbia (UBC).

## Author Contribution Statement

The study was conceived and designed by C.A.S, K.M.M, R.S., A.M.C. and A.C. Samples were collected by R.S. and S.W., A.S. and K.X.Z contributed to laboratory processing, sequencing and analysis steps. A.C. performed the lab work, A.C. and J.F.F. performed the bioinformatic and statistical analyses and interpretation with input from C.A.S., K.M.M., A.C. and K.X.Z. A.C. and J.F.F. wrote the manuscript with input from all co-authors.

## Conflict of Interest

The authors declare no conflict of interest.

## Supplemental Material file list

**Supplementary Table 1:** Pairwise Kruskal-Wallis test results comparing Shannon Index of the ‘low mortality’ and the ‘high mortality’ cohorts.

**Supplementary Table 2:** OTUs designated as the core microbiome with normalized abundance values per sample, taxonomic lineage per OTU is based on the SILVA database. Frequency is derived from the spat sample abundances, differential expression is derived from abundances in water and spat samples.

**Supplementary Table 3:** Linear models with R^2^, slope and significance (P-value) for the change in frequencies of significant core OTUs across age classes. Indval column describes OTUs that were determined significant in the indicator species analysis with ABC being early age classes and DEF being late age classes. Balance column describes OTUs that were included in the balance tree analysis of community composition versus age by samples, NUM in the numerator and DEN in the denominator of the balance calculation.

**Supplementary Table 4:** OTUs included in the neutral community model analysis with observed frequency and mean relative abundances in oyster samples (target communities), and water samples (source communities). The neutral model calculation includes the predicted frequency, the lower and upper 95 % limits, and the dispersion mode (Disp.). Neutral designates OTUs likely dispersed through the water column, Oyster designates OTUs specialized to oyster habitat and Water designates OTUs specialized to the water column.

**Supplementary Figure 1:** Constrained dbRDA of the community composition of oyster sample microbiomes against mortality. Blue icons indicate healthy samples with no abnormal mortality, red icons are samples with high mortality.

**Supplementary Figure 2:** Linear model of the calculated balance values and corresponding age of core OTUs, R^2^ of 0.9266 and a P-value of <2e-16.

## Notes

### Competing Interest Statement

The authors have declared no competing interest.

