## Supplementary figures and images for "The core microbiome of cultured Pacific oyster spat develops with age but not mortality"

### SupplementaryFigure1.pdf

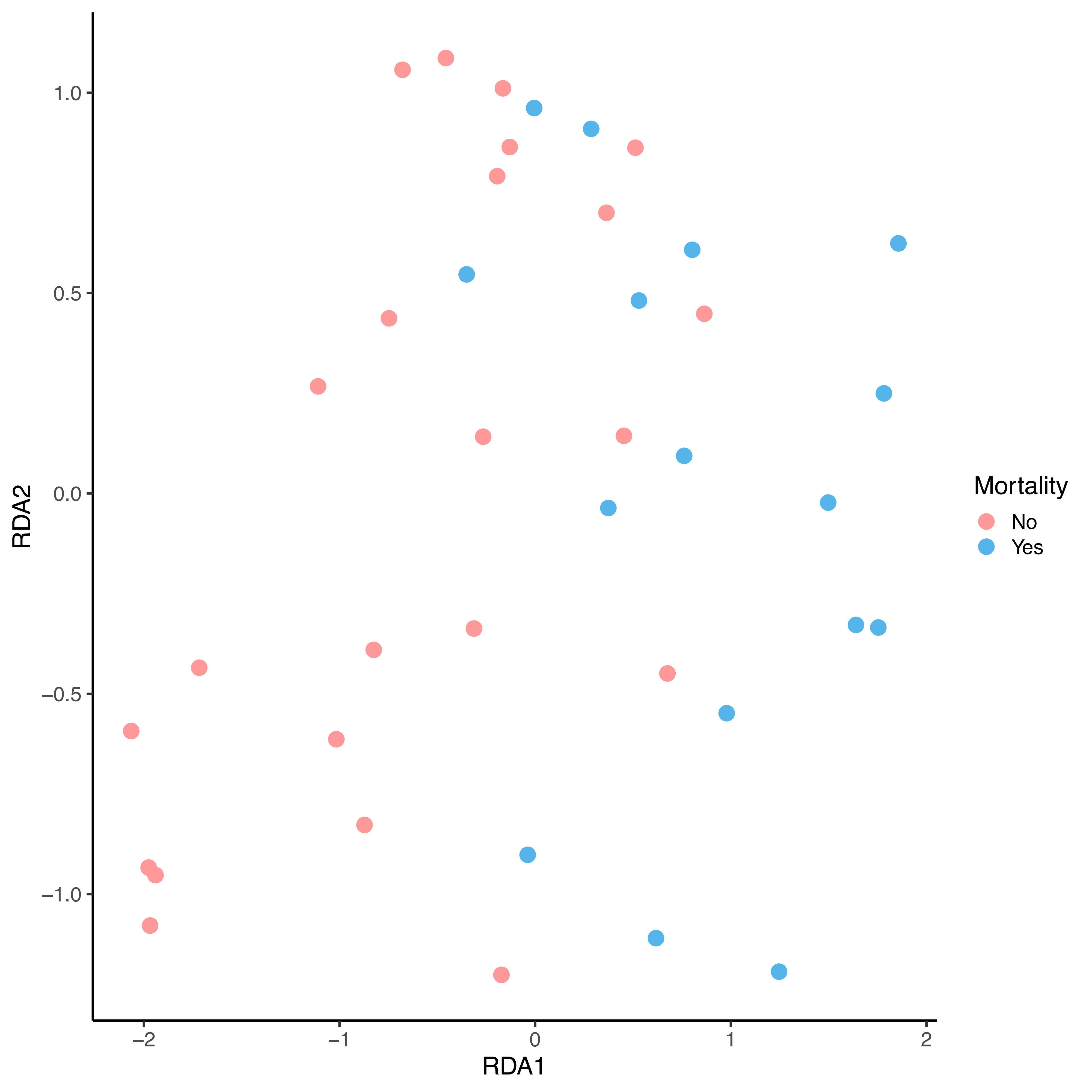

### SupplementaryFigure2.pdf

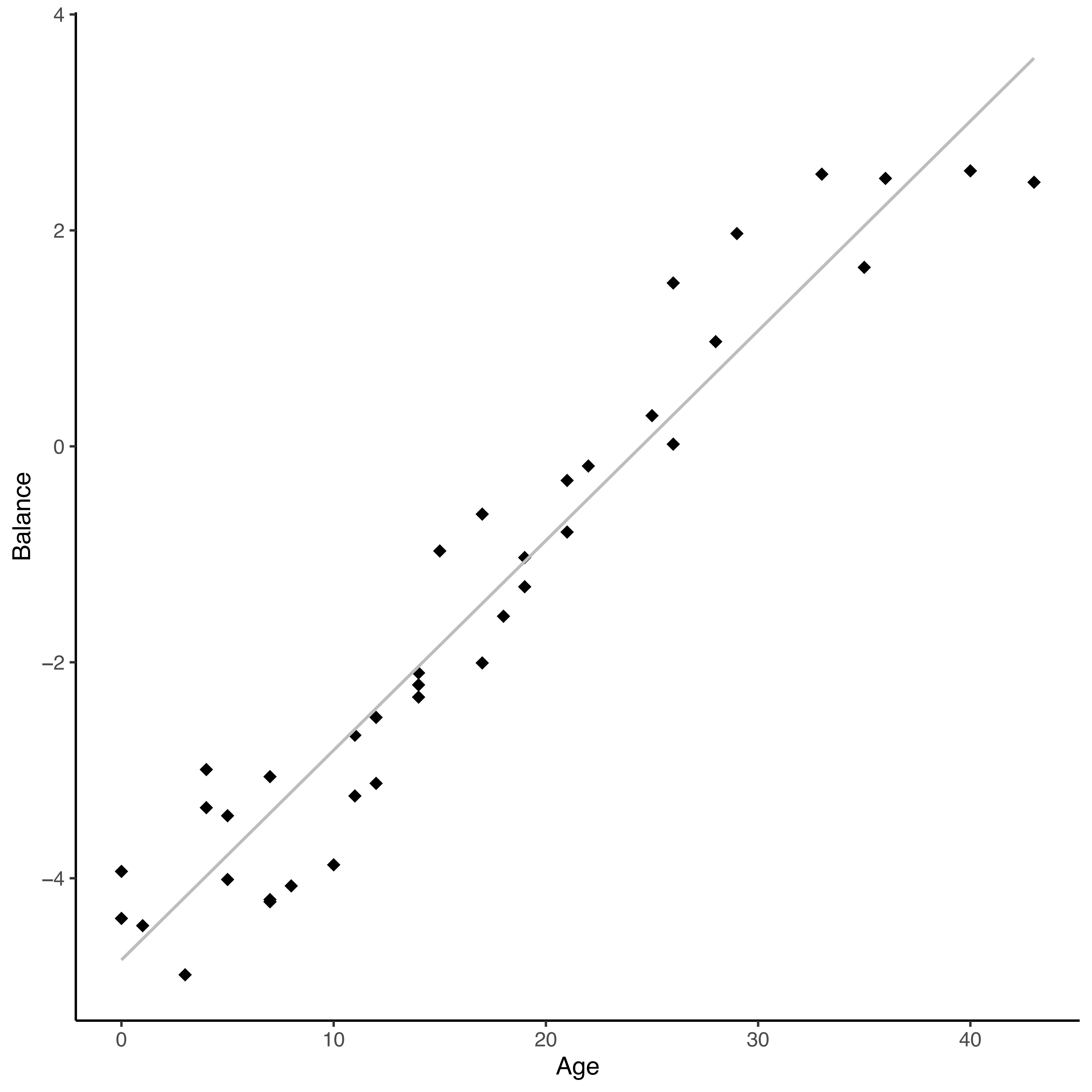
